# Applying a metaweb approach to reserve design: large, well protected areas are crucial to maintain food webs

**DOI:** 10.64898/2026.03.01.708826

**Authors:** Théo Villain, Hippolyte Erve-Sauvez, Jean-Christophe Poggiale, Charlie Marsily, Nicolas Loeuille

## Abstract

Establishing protected areas is a promising tool to address the accelerating loss of biodiversity. However, protection levels are often low, and there is an ongoing debate over the most effective spatial configuration of reserves. This debate rarely considers trophic structure and ignores biodiversity outside protected areas. In this study, we investigate which reserve configurations best support species diversity and the persistence of high trophic levels, across systems and spatial scales, both inside and outside protected areas. Using a spatially explicit stochastic model, we assess how reserve architecture influences multiple conservation objectives across 27 empirical terrestrial, freshwater, and marine food webs. Specifically, we explore reserve architecture along three dimensions: the aggregation of protected areas, their proportion at the landscape scale, and the effectiveness level of protection measures. Our results show that having few but larger protected areas enhances all conservation metrics within reserves, while α diversity within and outside reserves is relatively insensitive to reserve aggregation. Smaller and more dispersed reserves improve the overall abundance of species off-reserves through spillover effects. Reconciling all objectives inside and outside reserves becomes feasible when protection effectiveness is sufficiently high. Increasing the efficiency of protection allows for a reduction in the total amount of protected land without compromising conservation outcomes. Moreover, higher species dispersal facilitates the achievement of multiple conservation goals, supporting the implementation of architectures that enhance connectivity among reserves. These findings highlight the importance of an integrated approach combining spatial ecology and trophic functioning to optimize protected area planning under multiple objectives.

## Introduction

The impact of human activities on biodiversity is increasingly significant and is unfortunately driving the sixth mass extinction on Earth (Cowie et al., 2022; McCallum, 2015). Humanity relies directly or indirectly on the ecosystem services provided by biodiversity for food, energy, and resources (Fitter et al. 2010; Mace et al., 2012). The loss of biodiversity is thus a critical issue on its own, but it also poses significant threats to human activities (Dobson et al., 2006; Worm et al., 2006). To address this crisis, scientists advocate for a fundamental revision of humanity’s relationship with biodiversity, such as rethinking the management of fisheries (Pikitch et al., 2004), forests (Lindenmayer et al., 2000; The Montréal Process 2015), agricultural lands (Dudley and Alexander, 2017; Rey Benayas and Bullock, 2012), and the protection of some areas from human activities (Grorud-Colvert et al., 2021). Protected areas, both terrestrial and marine, have emerged over the past five decades as a key tool for conserving biodiversity. Evidences show that protection enhances species biomass and abundance (Lester et al., 2009; Soykan and Lewison, 2015), increases species diversity (Blowes et al., 2020; Gray et al., 2016; Lester et al., 2009), boosts ecosystem services (Sala et al., 2021; White et al., 2008) and improves the resilience of protected communities (Mellin et al., 2016), particularly in the face of climate change (Lehikoinen et al., 2019).

The effectiveness of protected areas first depends on pre-defined objectives. Designing protection to safeguard endangered species (e.g., Venter et al., 2014), particular communities (Mellin et al., 2016), or to maximize ecosystem services (Vandeperre et al., 2011) involves distinct approaches and implications. Protected areas may however support multiple objectives simultaneously, even when these may initially seem conflicting (Sala et al., 2021). For instance, protecting 28% of the world’s oceans could theoretically lead to the overall maintenance of biodiversity, to an increase of 6.2 million metric tons in fish catch and to the mitigation of carbon release associated with sediment disturbance caused by bottom trawling (Sala et al., 2021). Implementation of reserves however often generates social conflicts (Kovács et al., 2015) and constrain economic development in certain regions particularly because of short-term socio-economic costs (chapter 14 in Schreckenberg et al., 2018). Thus, while protection generally yields benefits across multiple dimensions, there are synergies and trade-offs between the objectives pursued (Davies et al., 2018; Kovács et al., 2015; Sala et al., 2021).

The effectiveness of protected areas also depends on the quality of the protection measures and on the financial resources available to support them (Watson et al., 2014). In 2024, 17.6% of terrestrial areas and 8.4% of marine and coastal areas were under protection (UNEP-WCMC and IUCN 2024). These figures mask significant disparities, as they include a wide range of protection levels —from strictly protected areas (where human presence is prohibited) to zones managed to optimize resource production (IUCN category VI). Many so-called “paper reserves” exist, where protection measures and enforcement are either absent or inadequate (Watson et al. 2014). According to Baillie and Zhang (2018), one-third of terrestrial protected areas are subject to intense human exploitation, and only 2.9% of marine areas are effectively protected (Dureuil et al., 2018; Grorud-Colvert et al., 2021; MPAtlas 2024).

The effectiveness of protection also depends on the spatial design of reserves. There is a broad consensus on the need to increase the total area of the planet under protection to safeguard a greater diversity of ecosystems and species (Allan et al., 2022; Baillie and Zhang, 2018; Larsen et al., 2015; Visconti et al., 2019). Among the various possible spatial configurations of reserves, those that best protect biodiversity are the ones where the total amount of protected habitat, regardless of its configuration, is the largest (*habitat amount hypothesis*, Fahrig, 2013). These calls have driven the transition from the Aichi Biodiversity Targets set in 2011 (17% of terrestrial and 10% of marine areas, CBD 2011) to the more ambitious goal of protecting 30% of all ecosystems (both terrestrial and marine) by 2030, as outlined in Target 3 of the Kunming-Montreal Global Biodiversity Framework. However, the emphasis on expanding the overall protected area should not be conflated with the size of individual reserves. Within each protected area, the surface under protection remains a key determinant of conservation success (Claudet et al., 2008; Gaines et al., 2010), especially for highly mobile species (Boersma and Parrish, 1999). Nonetheless, the specific effects of reserve size (whether additive or amplifying) vary depending on the ecological and socio-economic context (Halpern, 2003; Maiorano et al., 2008).

Once the total reserve size is set, an important question is how this total protected area should be set in space. Depending on underlying objectives, it may be better to create few large or many small protected areas (the SLOSS debate, Single Large or Several Small, e.g., see Fahrig et al., 2022). At a broader scale, the effectiveness of a reserve in conservation depends on the presence of nearby reserves and, consequently, its connectivity (Brennan et al., 2022, Gaines et al., 2010). For the same total area under protection, there is emerging empirical evidence favoring such a “*several small*” strategy: the greater the number of distinct reserves, the higher the species richness reported across the entire network of reserves (e.g., reviewed by Fahrig, 2020). At this stage, it remains challenging to determine whether these results stem from a higher probability of encompassing a broader range of habitats - and therefore distinct species assemblages - when multiple reserves are implemented, or whether they truly result from the spatial configuration of the reserves themselves. Indeed, species richness patterns across multiple reserves may reflect both habitat heterogeneity and vertical diversity (Scheiner et al., 2011).

The SLOSS debate most often focuses on the maintenance of species richness, regardless of the structure of ecological networks. It is however well-known that network aspects (eg, being a specialist or at higher trophic levels) largely constrain the vulnerability of natural populations. These approaches also rarely account for the role of naturality in areas outside reserves. Beyond its intrinsic ecological value, naturality can profoundly affect both human well-being and the delivery of ecosystem services. For instance, marine protected areas have been shown to boost fishery yields in surrounding waters (Sala et al., 2021; Vandeperre et al., 2011), when adjacent exploited zones benefit from the spillover of adult individuals (Gaines et al., 2010). In terrestrial ecosystems, comparable spillover effects can have key functional implications (Blitzer et al., 2012), such as enhancing pollination in agricultural landscapes (Garibaldi et al., 2011) or facilitating pest control through predation (Rusch et al., 2010). These economic implications of off-reserve naturality come in addition to the intrinsic value of having healthy ecosystems on the overall landscape, not just in protected areas.

While many theoretical studies have explored spatial aspects of reserve design, they most often overlook the complexity of underlying ecological networks. Most modeling work remains focused on single-species dynamics (for a review, see (Burkey, 1989; McCarthy et al., 2005; Ovaskainen, 2002) or on competition-based metacommunities (Chase et al., 2020; Tjørve, 2010), with subsequent extrapolations to network-level outcomes (Fahrig, 2020). Trophic interactions are rarely integrated (McCarthy et al., 2006; Pardini et al., 2010; Tjørve, 2010), despite it could significantly shift our understanding of reserve effectiveness. For instance, the maintenance of upper trophic levels relies on prey availability (Gravel et al 2011), so that we expect that efficient reserve designs for upper trophic levels are likely a constrained subset of designs applicable for lower levels. Consistently, the impact of habitat fragmentation may vary depending on a species’ trophic position (Rielly-Carroll and Freestone, 2017) and on the time elapsed since reserve establishment (Claudet et al., 2008; Soulé and Simberloff, 1986; Vandeperre et al., 2011). Furthermore, the trophic structure of ecological networks is fundamental to their stability (Landi et al., 2018) and resilience to species loss (Keyes et al., 2024). While connectivity between reserves consistently enhances persistence in metapopulations, it has contrasting effects in metacommunities based on competition (e.g., Mouquet and Loreau, 2003). In metawebs, connectivity could modulate total and vertical diversity in different ways depending on the dominant control regime (top-down vs. bottom-up).

To address these limitations, we modeled the impact of reserve configuration on the spatial persistence of trophic networks. We try to understand how reserve design contributes to preserving local and global diversity at landscape scale, inside reserves but also how it preserves naturality outside through spillover effects. Inspired by the rationale of the *trophic theory of island biogeography* (Gravel et al., 2011), we have developed a spatially explicit stochastic model where complex ecological networks are considered to test different reserve designs using a multi-objective approach. Our evaluation employs metrics of diversity and integrity, including species richness and ecosystem verticality, and compares a broad array of terrestrial and aquatic networks from various databases. Specifically, we seek to:

1. Contribute to the SLOSS debate through a network-based perspective.
2. Identify reserve configurations (aggregation of reserves, proportional coverage of the landscape and level of protection) that sustain ecosystem functionality while simultaneously delivering ecosystem services outside reserves.
3. Understand whether it is possible to substitute one axis for another and achieve the same conservation status.

We hypothesize that higher levels of protection enhance the multi-dimensional effectiveness of conservation measures. The proportion of protected area is expected to enhance ecosystem services when spillover effects are significant, provided that the remaining exploitable area is sufficiently extensive to benefit from these inputs. For spatial aggregation, our predictions are more nuanced. On one hand, by creating large, high-quality areas with complete networks, aggregation can increase fluxes across reserve boundaries, thereby promoting ecosystem services in adjacent exploited zones. On the other hand, by reducing the edge-to-area ratio, it may limit the extent of interfaces where spillover can occur.

## Methods

### 1) Protected area architecture

In this study, we aim to identify the optimal reserve configuration to meet different conservation objectives across various spatial scales. We consider a landscape modeled as a 50×50 grid of patches, structured as a torus to avoid edge effects (Figure 1). Within this landscape, areas can either be protected (PAs, colored pixels in Figure 1) or exploited in the sense that higher levels of human activities are allowed (EAs, white pixels). This design is investigated along three axes: (i) the proportion of the grid that is protected (i.e., the total surface of PAs), (ii) the spatial arrangement of the PAs, and (iii) their level of protection (Figure 1). We explore a wide range of protection proportions (from 0% to 100% of the landscape; “proportion” axis in Figure 1), ranging from random to spatially autocorrelated reserve layouts (“aggregation” axis in Figure 1), and with different protection efficiencies depending on the scenario (“protection” axis in Figure 1; *p* = 0. 2 for weak protection in black, *p* = 0. 3 for intermediate protection in light green, and *p* = 0. 5 for strong protection in dark green). To generate different levels of spatial aggregation, we use six spatial autocorrelation indexes (SAI) with the *mdp* function from the NLMpy package (Etherington et al., 2015), spanning from highly dispersed to highly aggregated configurations (*SAI* = −10, −1, 0, 0.5, 1, or 2). These various reserve architectures are then tested across 27 empirical trophic networks.

**Figure 1:**
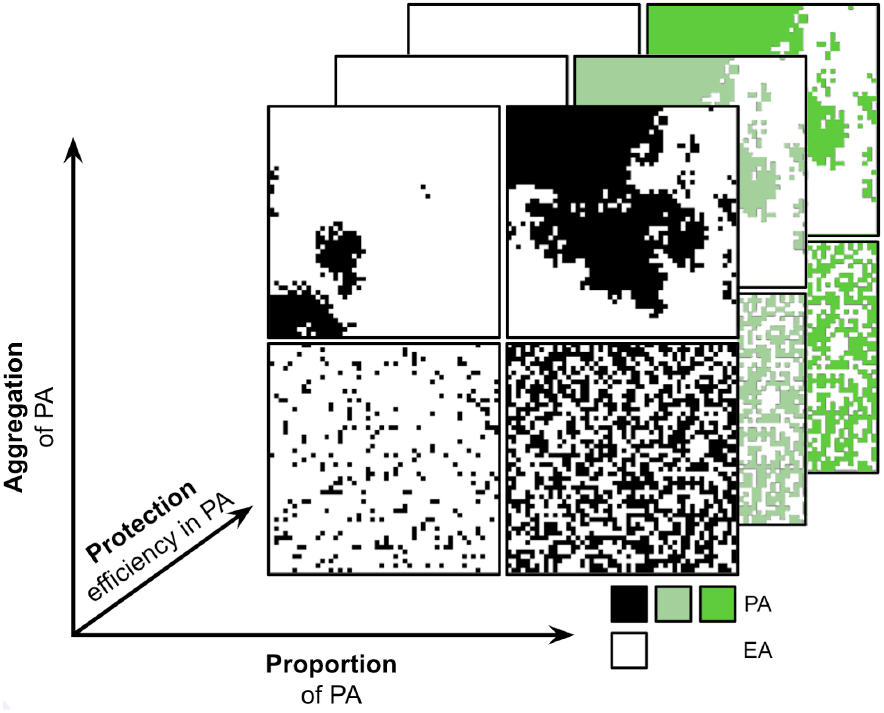
Design of protected area configurations varying in spatial aggregation, landscape coverage, and protection level. Colors indicate the degree of protection. PA: Protected Area; EA: Exploited Area.

### 2) Ecological model and spatial dynamics

To understand the influence of reserve configurations, we developed a model that incorporates trophic interactions among species within the same network and embeds them in a spatially heterogeneous protection context. The model builds on the assumptions of the *trophic theory of island biogeography* (Gravel et al., 2011), which relies on extinction-colonisation processes in which predators only colonize and survive if prey species are locally present. To fit our questions, we modify the model in two ways: (*i*) colonization-extinction also depends on whether a given area is protected or exploited; (*ii*) we account for top-down and bottom-up forces in the sense that colonisation-extinction for a given species depends not only on prey (as in the original model), but also on predator presence. These processes are unified and made continuous by defining a local quality *Q*_*i,j*_ of a patch *j*, specific to the dynamics of species *i* (eq. 1).

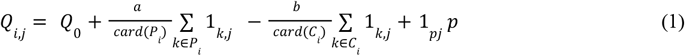

Patch quality depends on a baseline value *Q*_0_, on the presence of prey (with *P*_*i*_ denoting the set of allpotential prey for species *i*), of predators (with *C*_*i*_ denoting the set of all potential predators), and on whether the area is protected or exploited (with 1_*pj*_ equal to 1 when the patch is protected and 0 otherwise). Locally, only a subset of prey and predator populations may be present, which is accounted for by indicator functions 1_*K,j*_ (equal to 1 when species k is present, 0 otherwise, with *K* ∈ *P*_*i*_ or *K* ∈ *C*_*i*_). For species that have no predators in the ecological network, the third term in (1) is omitted. The model thus considers the presence or absence of species rather than their abundances. Parameter *a* represents the contribution of prey to patch quality, while *b* reflects its degradation by predators, these two parameters thereby reflecting the relative weight of bottom-up vs top-down constraints. A more complete the predation regime for species *i* therefore leads to higher patch quality (Figure 2*B*_*j*_ - *B*_*j*′_-C). Conversely, a more complete predator set locally reduces patch quality (Figure 2 *B*_*j*_ - *B*_*j*′_-C). For a given species composition, protection level *p* enhances patch quality (eq. 1 and Figure 2C).

**Figure 2:**
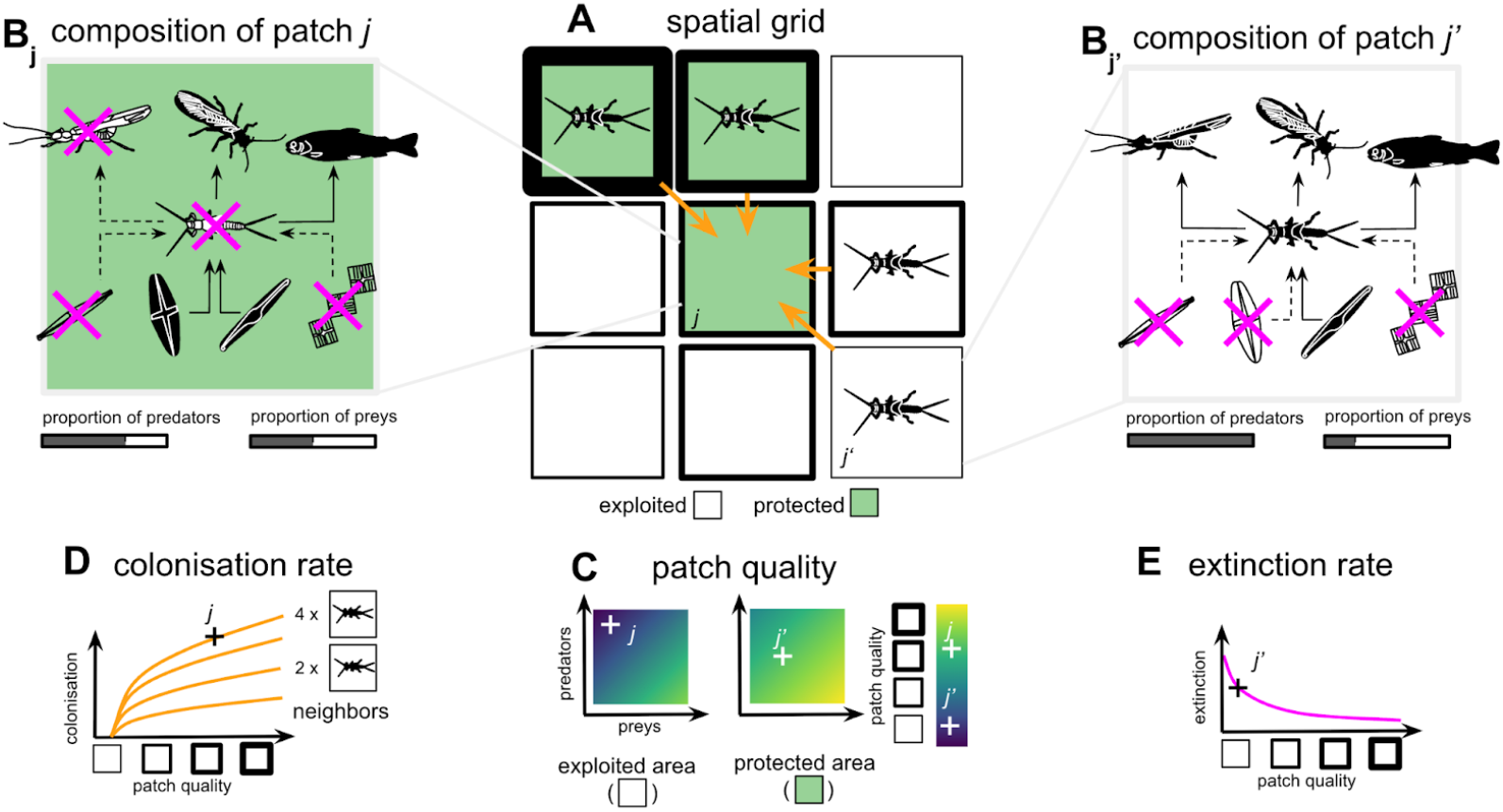
Patch quality and spatial dynamics associated with colonisation-extinction processes. A: Subsection of the grid; the focal species is present in 4 out of 9 patches. Green patches: protected; white patches: exploited. Patch border thickness reflects the quality of the patch for the focal species. B: Species composition in patch j (*B*_*j*_) or patch j′ (*B*_*j* ′_ here of lower quality, as it is not protected, with a low-prey high-predation context) among species interacting with the focal species. The focal species is absent from patch j, which can thus be colonized, and present in patch j′, where it can go extinct. C: Patch quality for the focal species as a function of prey proportion, predator proportion, and protection status. Patch quality determines the probability of colonization (D) and extinction (E) events.

The probability *c*_*i,j*_ that species *i* colonizes patch *j* and successfully establishes increases with the patch quality *Q*_*i,j*_, the proportion *n*_*i,j*_ of neighboring patches within its dispersal range that are occupied by species *i*, and the baseline colonisation probability *c*_o_ (eq. 2 and Figure 2D):

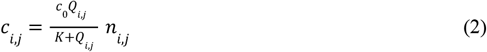

The dispersal distance is set to 1 for all species within a given network, meaning that only the 8 neighboring patches surrounding the focal patch are considered (Figure 2A). An additional analysis is performed using larger dispersal distances (Figures S4 and S5). *K* represents the half-saturation constant (the quality at which colonisation has half its potential max), and *c*_0_ corresponds to the maximum colonisation probability when the patch quality is infinite and all neighboring patches are occupied. A species *i* present in patch *j* can go locally extinct with a probability *e*_*i,j*_ decreasing with patch quality *Q*_*i,j*_ (eq. 3 and Figure 2E):

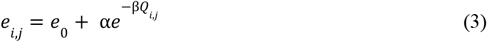

In high quality patches, the extinction probability tends toward a low value *e*_0_ (eq. 3). Parameter α represents the weight of the patch quality function on extinction probabilities, and β denotes the sensitivity of extinction probabilities to patch quality variations. In practice, both *c*_*i,j*_ and *e*_*i,j*_ are bounded between 0 and 1. When a random draw from a uniform distribution exceeds one of these probabilities, a colonisation or extinction event is considered to occur. For primary producers, it is assumed that all their resources are naturally present on the patches (resource constraint *a* is identical and uniform). Note that because prey is only part of the definition of quality, predators can survive for a slightly longer time than the strict disappearance of their prey, thus relaxing the concept of bottom-up sequential dependency (Gravel et al., 2011). Note however that our basic scenario considers that quality is more affected by prey than by predator presence (*a*>*b*).

To simulate the spatial dynamics of a trophic network within a given reserve configuration, the grid is first randomly seeded with the different species of a specific network (with a probability of 0.5 for each species). At each time step, the dynamics are analyzed patch by patch. For each species, the quality of the patches is determined based on the presence of their prey, their predators, and the protection status of the patch. Colonisation and extinction events are then drawn accordingly. Equilibrium is assumed to be reached at *t* = 500 at which point the fulfillment of the various conservation objectives is quantified. For each reserve configuration and each network, 10 replicates are performed.

### 3) Trophic networks

We apply this model to 27 different empirical trophic networks (2 terrestrial networks, 6 marine networks, and 19 freshwater networks). These networks were selected to cover a wide range of structures (species richness, connectance, maximum and mean trophic level) and ecosystem types. They were sourced from the Interaction Web Database (https://www.nceas.ucsb.edu/interactionweb/), the Global Food Web Database (https://www.globalwebdb.com/), and the R package *cheddar*. Details and characteristics of the networks used are provided in Figure S1 in the Supplementary.

### 4) Conservation objectives

To understand which reserve configurations are most suitable for maintaining biodiversity and ecosystem service provision, we use metrics at both local (mean patch-level properties within a set) and regional scales (properties of the entire set of patches). These metrics can be calculated for the whole landscape (PA + EA), only within protected areas (PA), or only within exploited areas (EA). At the local scale, we measure average α diversity (eq. 4) and network verticality (maximum trophic level within a patch, eq. 6). At the regional scale, we measure γ diversity, the verticality of the maintained regional network (maximum global trophic level, eq. 7), and off-reserve naturality (ORN, eq. 8), which quantifies the total population number outside protected areas.

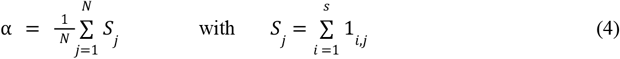

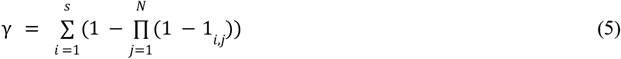

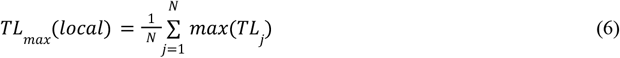

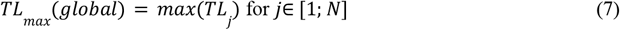

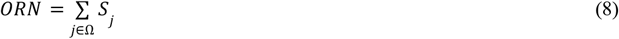

*N* is the number of patches in the studied set (PA, EA or EA+PA), *s* is the number of species in the full network, 1_*i,j*_ is the indicator function of the presence of species *i* in patch *j* (=1 if the species is present, 0 otherwise), *TL*_*j*_ is the set of extent trophic levels in patch *j*, and Ω is the set of exploited areas.

Since the establishment of protected areas is based on quantitative targets, a spatial configuration is considered effective in achieving a given objective if the measure of the associated property exceeds a certain threshold. When 35% of the local diversity of a network is maintained (α diversity), it is considered adequately protected with respect to this objective. This threshold is set at 80% for regional diversity (γ), 50% for the local maximum trophic level (*TL*_*max*_ (*local*)), 80% for the globalmaximum trophic level (*TL*_*max*_ (*global*)), and 5% for naturality outside protected areas (ORN). While these thresholds are arguably somewhat arbitrary and chosen to properly contrast outcomes in the result section (eg, figures 4 and 5), our analysis will focus on the qualitative effects of reserve architectures.

To highlight and explain the effect of spatial configuration on temporal dynamics and persistence, we first focus on a specific network (Afon Hafren, a stream ecosystem that has low species richness, high connectance and low verticality). In the second part, we investigate which reserve architectures are best suited to various conservation objectives. Finally, we examine which configuration allows the reconciliation of these different objectives.

**Table 1:** Variables and parameters used in the model.

| Index | Meaning | Value |
| --- | --- | --- |
| $i$ | species | - |
| $j$ | patch | - |
| Functions | Ecological meaning | Value |
| $Q_{ij}$ | Quality of patch $j$ for species $i$ | [0;1] |
| $c_{ij}$ | Colonisation probability | [0;1] |
| $e_{ij}$ | Extinction probability | [0;1] |
| $1_{ij}$ | Indicator function of species $i$ presence in patch $j$ | 0 (absent) or 1 (present) |
| $1_{pj}$ | Indicator function of patch $j$ protection | 0 (exploited) or 1 (protected) |
| Variables | Ecological Meaning | Value |
| $P_i$ | Set of prey species which interact with $i$ | {} |
| $C_i$ | Set of predator species which interact with $i$ | {} |
| Parameters | Ecological meaning |  |
| $c_0$ | Maximal colonisation intensity | 3 |
| $e_0$ | Basal extinction intensity | 0.01 |
| $Q_0$ | Basal patch quality | 0.05 |
| $n_{ij}$ | Patch proportion within the dispersal area around patch $j$ where species $i$ is present | [0;1] |
| $a$ | Prey contribution to species $i$ patch quality | 0.4 |
| $b$ | Predator degradation of species $i$ patch quality | 0.15 |
| K | Half-saturation constant for colonisation probability | 2 |
| $\alpha$ | Weight of the quality function on extinction | 0.9 |
| $\beta$ | Sensitivity of extinction to patch quality | 2 |
| p | Protection effect | {0.2,0.3,0.5} |

## Results

### 1) Spatial configuration and temporal dynamics for a specific network

Regardless of the reserve architecture, the first species to disappear are those at high trophic levels (Figure 3). These species have a lower probability to find and colonize patches where their prey are present, as these prey depend on the presence of their own resources. The proportion of protected areas and their aggregation promote the conservation of various species, particularly at the landscape scale (red and purple curves). Local diversity measures (α diversity and local maximum trophic level) as well as off-reserve naturality stabilize quickly at low levels (between 15% and 50%) and are here little affected by reserve architecture. For this particular (freshwater) network, aggregation of patches allows maintaining a complete network at the landscape scale (insets in Figures 3A and 3B), while locally only 20 to 25% of species are present (Figures 3A and 3B). The worst landscape configuration is that with few reserves distant from each other (Figure 3C), where all top predators and herbivores rapidly disappear.

**Figure 3.**
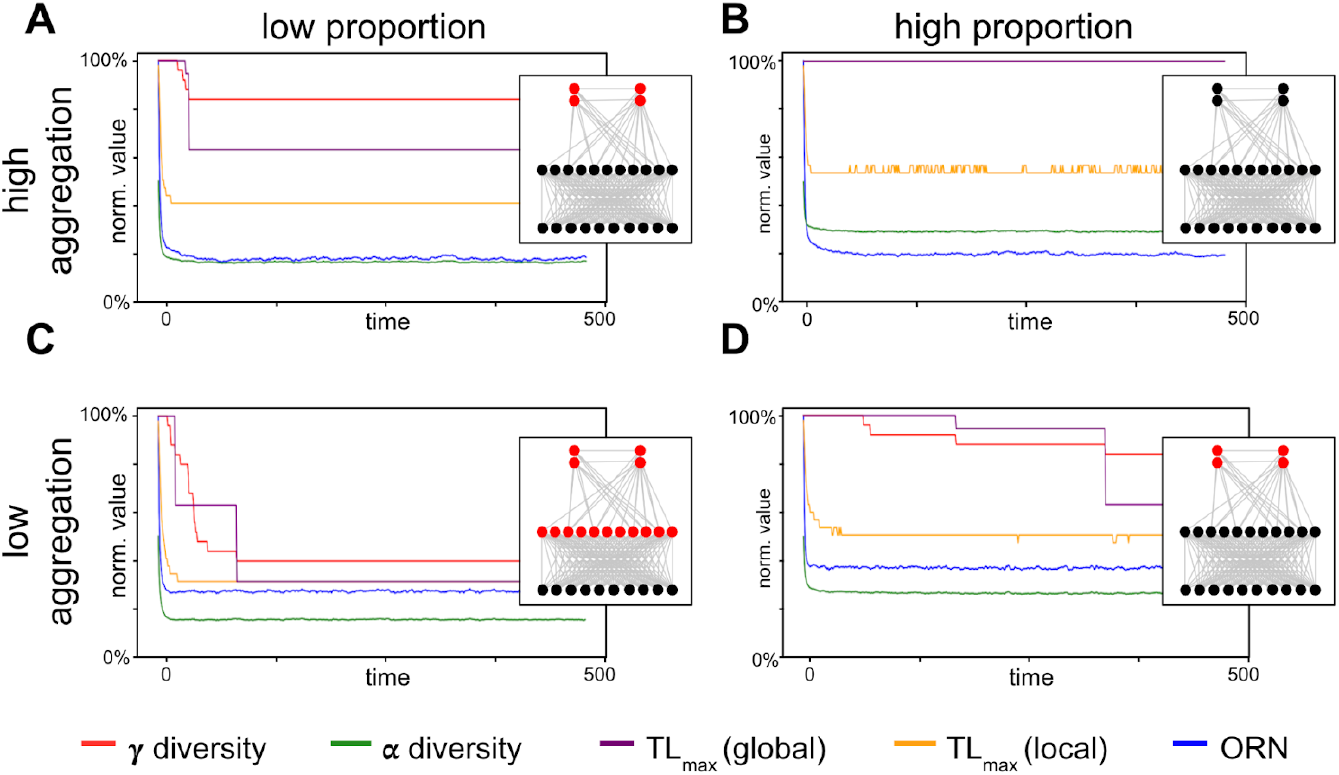
Effects of aggregation and proportion of protected areas on the composition of the Afon Hafren trophic network. Within the landscape, protected areas (PA) can be highly aggregated (A and B) or weakly aggregated (C and D). The proportion of PA can be high (B and D) or low (A and C). After random seeding of the grid, species colonize or go extinct in patches depending on patch quality. These processes affect γ diversity (red), α diversity (green), maximum trophic level across the entire grid (purple), maximum trophic level per patch (yellow), and off-reserve naturality (blue). At equilibrium (*t* = 500), a representative trophic network of the entire landscape is shown, extent species being in black and extinct species in red (insets). The protection level within PA is intermediate.

### 2) Considering all networks, large reserves protect diversity at the landscape scale but penalize off-reserve naturality

Large, highly aggregated reserves are optimal for meeting diversity objectives (local, global and verticality) at the landscape scale (Figure 4A–I), whereas medium-sized, dispersed reserves maximize naturality in exploited areas (Figure 4J–L). Regardless of spatial configuration, higher protection levels consistently enhance the fulfillment of conservation objectives.

**Figure 4:**
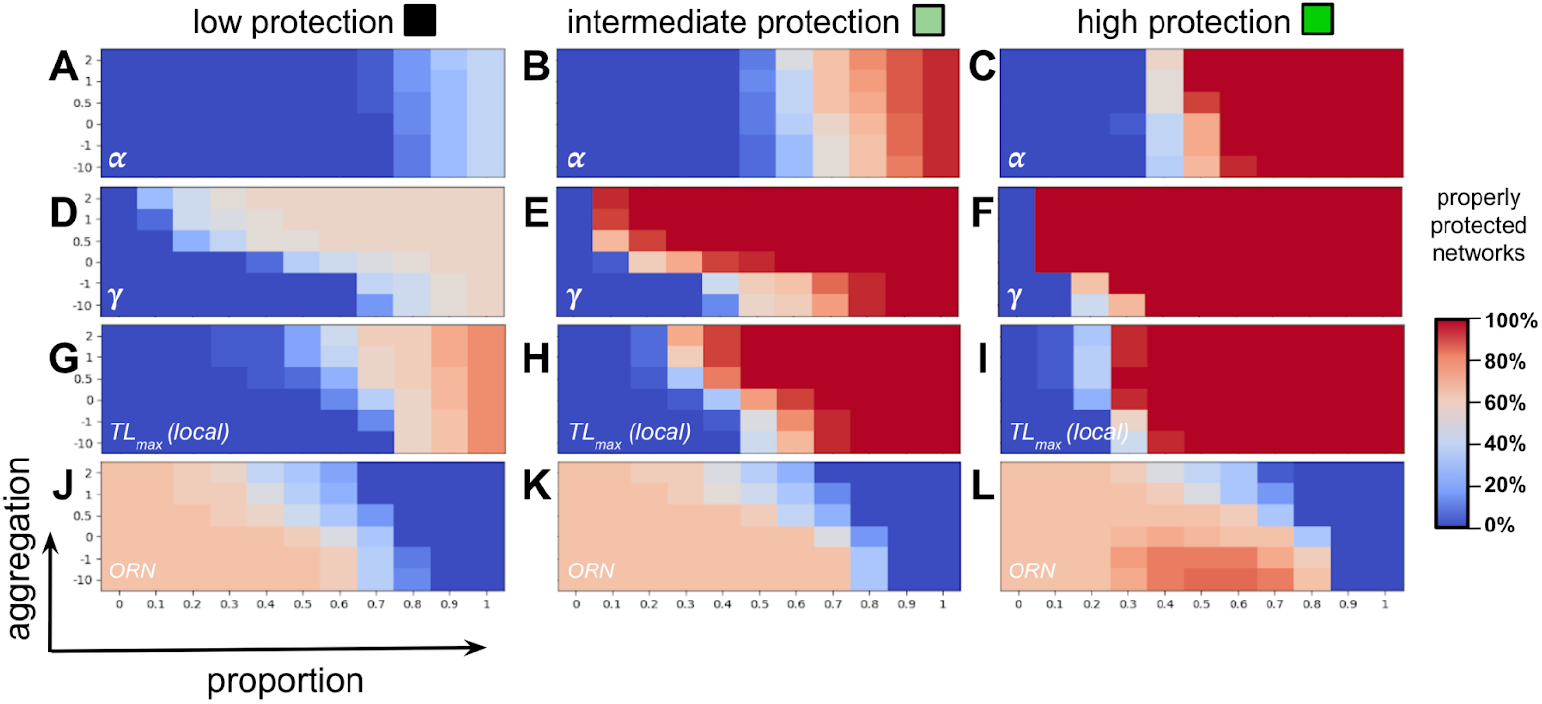
Effectiveness of reserve architecture (aggregation, proportion, and protection level) on achieving conservation targets across 27 different networks. A spatial configuration is considered effective for a conservation objective when it maintains the network state above the corresponding threshold. When the network maintains at least 35% of its local diversity (α, A, B, C), it is considered adequately protected. This threshold equals 80% for regional diversity (γ, D, E, F), 50% for the maximum local trophic level (G, H, I), and 5% for naturality outside reserves (ORN, J, K, L). The three columns differ by protection level, which are, from left to right, *p* = 0. 2, *p* = 0. 3 and *p* = 0. 5.

Interestingly, the management of diversity is however slightly different depending on the spatial scale. Locally (alpha diversity), a suitable protection mostly depends on the proportion of landscape that is protected, aggregation only slightly changing the results. At a larger scale (γ diversity, Figures 4D–F), high global diversity can be obtained for smaller proportions, if protected areas are sufficiently aggregated. Increasing aggregation, proportion, and protection level leads to reserves that are more numerous, more contiguous, and of higher quality, which promotes the maintenance of highly diverse patches within them and thus meets the conservation objective (Figure S1D–F). These parameters also support conservation in exploited areas (Figure S3D–F), as they allow for the presence of diverse patches at the edges of protected areas.

Effects of reserve architecture on naturality off reserve (and possible ecosystem services it sustains) are vastly different (figure 4J-L). First, because naturality is computed using the total number of populations present in exploited patches, increased proportion of protected patches has a dual effect. First, increased proportion of protected patches mathematically decreases the maximum number of populations that are off-reserves (negative effect on naturality). Second, higher proportion of reserves feed adjacent areas through spillover effects (positive effect on naturality). Because spillover is more efficient when the surface of contact is important, off-reserve naturality is better managed using non-aggregated landscapes, at intermediate proportion of protection (eg, figure 4L).

Aggregation however creates large areas devoid of reserves, and low reserve proportion limits the number of exploited patches. This explains why the effect of aggregation is much less pronounced on landscape-scale α diversity (Figure 4A–C). In exploited areas, it is the dispersion of reserves that promotes α diversity (Figure S3A–C) through spillover effects, although diversity levels remain low (at best, 30% of networks are adequately protected). Thus, when the proportion of reserves is low, aggregation has little influence at the landscape scale (diversifying exploited areas requires dispersed reserves). However, as the proportion of reserves increases, aggregation exerts a stronger positive effect: protected areas become more diverse when they are aggregated (Figure S1A–C). These effects also explain why off-reserve naturality is highest with low aggregation and intermediate reserve proportion (Figure 4J–L). Increasing the proportion of reserves leads to more numerous and larger reserves, thereby generating stronger spillover effects into exploited areas. Conversely, the amount of exploited area decreases, hence the potential number of populations present in exploited areas. The total number of populations across the landscape peaks around 50-60% reserve coverage. Finally, local trophic verticality across the landscape (*TL*_*max*_(*local*), Figure 4G–I) follows similar dynamics, but the conservation target is more easily reached than for α diversity, notably because subnetworks are maintained within exploited areas (Figure S3G–I), allowing for the persistence of trophic chains.

Protection level can lead to abrupt changes in the global conservation status of food webs (Figure 4F). Increasing the aggregation of just a few highly protected reserves (10% of the patches) causes the number of well-managed networks (with respect to γ diversity) to jump from 0% to 100%, while it would have no effect if protection levels are low (Figure 4A). This highlights the importance of a minimal threshold size of high-quality reserves, as it ensures the persistence of highly favorable patches for diversity somewhere in the landscape. Finally, species dispersal has a similar effect to protection level and promotes the conservation of the various networks (Figures S4 and S5). The greater the mobility, the weaker the effect of reserve dispersion, since species can colonize patches located further from their original positions.

### 3) Stronger protection allows reducing the proportion of reserves in the landscape and concentrating them

We now assess the possibility of conciliation of the different objectives, by highlighting scenarios in which all objectives are met (Figure 5A-C). Because off-reserve naturality and diversity objectives require different spatial strategies (summarized on Figure 5D), conciliation is hard (panels A-C are mostly blue). Particularly, it is absent if protection is low (Figure 4A) and seldom occurs at intermediate levels (Figure 4C). As the level of protection increases, reconciling conservation objectives (local diversity, global diversity, local maximum trophic level, and off-reserve naturality) becomes easier and allows for fewer, less dispersed reserves (Figure 5).

**Figure 5:**
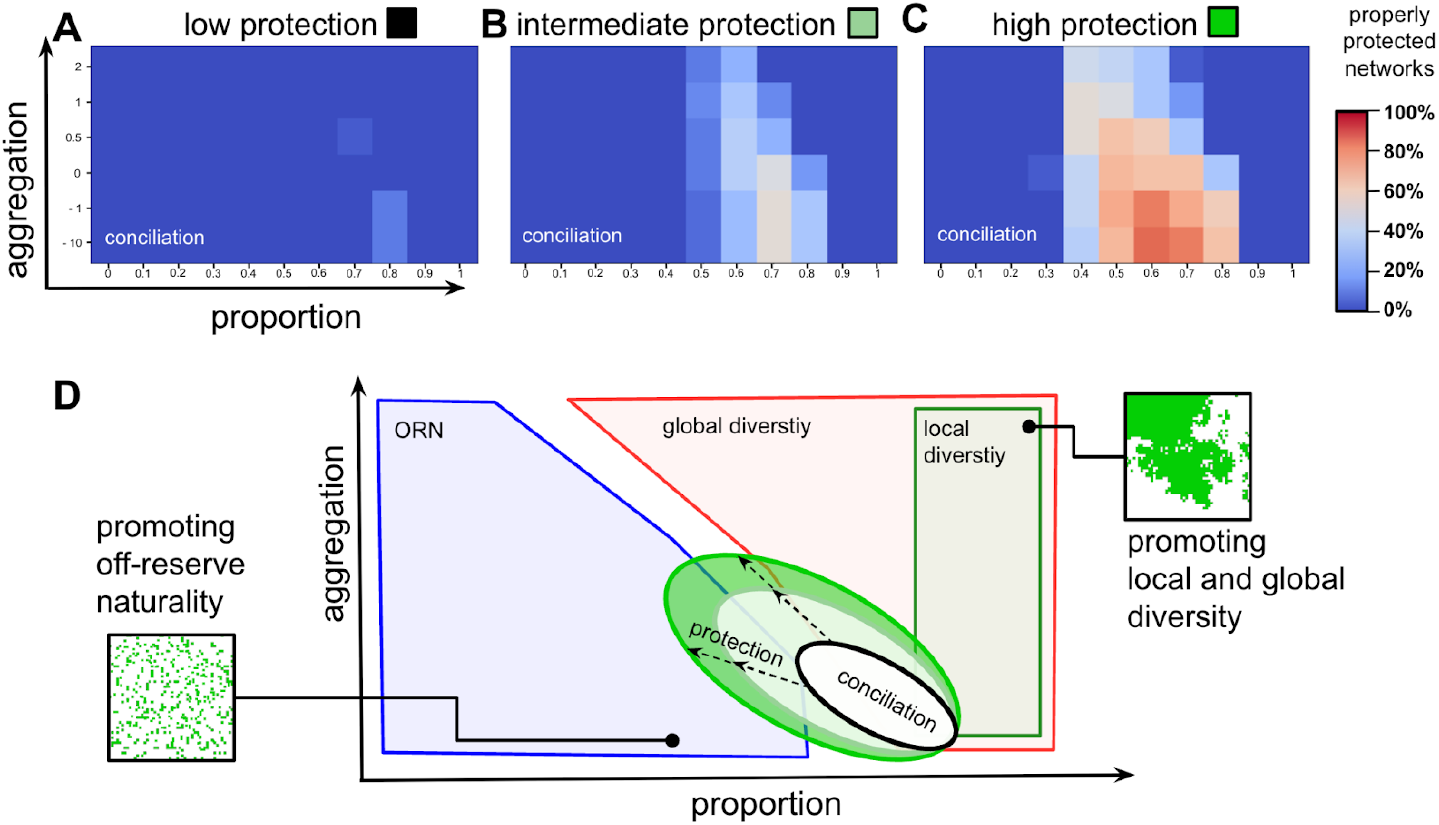
Reconciling multiple conservation objectives as a function of reserve architecture (aggregation, proportion, and protection) across 27 different food webs. A spatial configuration is considered effective for a given conservation objective when it maintains the state of the network above the corresponding threshold. A network is considered properly protected when it maintains at least 35% of its local diversity (α diversity). This threshold is set at 80% for global diversity (γ diversity), 50% for the local maximum trophic level, and 5% for off-reserve naturality. A–C: the protection level is respectively *p* = 0. 2, *p* = 0. 3 and *p* = 0. 5. D: Summary diagram showing the effectiveness of reserve architecture with respect to different conservation objectives.

Reconciling the various conservation objectives is challenging when protection levels are low (Figure 5A, *p* = 0. 2). When reserves are numerous (80%) but spatially dispersed, only 20% of food webs can be protected in a way that satisfies all objectives. As the level of protection increases (Figure 5B, *p* = 0. 3), a greater number of spatial configurations become suitable, including those with fewer and more aggregated reserves, but the proportion of well managed food webs remains low. Only intermediate levels of aggregation (spatial autocorrelation = 0) and 70% of the landscape under protection allow 50% of networks to be adequately conserved. Finally, when protection is even stronger (Figure 5C, *p* = 0. 5), up to 85% of food webs can be effectively protected with an even lower proportion of reserves (60%). These results clearly highlight the importance of suitable enforcement of protection to allow an integrative management of systems.. Protected areas then tend to become persistent source zones, and spillover effects are stronger, sustaining high diversity in exploited zones. More aggregation then becomes compatible with high off-reserve naturality, as surrounding protected areas are more species-rich. As exploited zones become more diverse, they provide higher-quality habitat for top predators, enabling stepwise colonisation of areas farther from reserves. However, when reserve coverage exceeds 90% of the landscape, it becomes impossible to reconcile all conservation objectives regardless of the protection level, as too few exploited patches remain to support significant biodiversity in it (Figures 5A–C and 4J–L).

## Discussion

Our study shows that effectiveness of reserve architecture depends on the conservation objectives and the spatial scale at which they are assessed. At the landscape scale, promoting local diversity requires a high number of reserves, while their aggregation matters little (Figure 5D, green box). This result stems from the fact that exploited areas typically support low diversity, which increases when reserves are spatially dispersed due to spillover effects. Conversely, local diversity is higher within protected zones when these are spatially aggregated. Aggregation has a stronger effect on the local maintenance of food web verticality, as highly aggregated reserves can sustain subsets of species that form trophic chains. Promoting global diversity at the landscape scale (Figure 5D, red) requires reserve aggregation, as it ensures the persistence of highly diverse zones. These configurations are effective both when considering diversity within protected areas and when assessing diversity in exploited zones, since patches near reserves benefit from significant spillover. To enhance off-reserve naturality, an intermediate number of rather dispersed reserves is needed (Figure 5D, blue). As species dispersal ability increases, reserve effectiveness improves across all conservation goals, by amplifying spillover effects. Finally, to reconcile these different objectives, a high proportion of reserves that are relatively dispersed is needed (Figure 5D). Our results also highlight that protection level is crucial, as high reserve effectiveness is highly important regardless of objectives, and essential for conciliation. As it increases, it becomes possible to conserve more networks with fewer, potentially more aggregated, reserves (Figure 5D, green circles).

### Protecting large surfaces

In response to accelerating biodiversity loss (Butchart et al., 2010), the Kunming-Montreal Global Biodiversity Framework recommends protecting 30% of all ecosystems by 2030. While extrapolating across scales remains contentious, our results indicate that this target is insufficient to sustain high levels of biodiversity across an entire landscape. At this proportion, trophic networks persist primarily within protected areas, and only under sufficiently high protection levels. Locally, exploited zones exhibit low diversity, and only 60% of trophic networks show off-reserve naturality exceeding 5% of their potential maximum. Our findings suggest that protection thresholds exceeding 50% of surface area would be needed to ensure biodiversity is maintained across spatial scales, to preserve trophic network functionality, and to allow exploited areas to benefit from biodiversity spillover (assuming that protection level is at least intermediate). At the reserve scale, this aligns with recommendations by Boersma & Parrish (1999), who showed that very large areas are required to protect wide-ranging marine species. Larger reserves are more effective at safeguarding species vulnerable to exploitation—such as commercial fish—than smaller ones (Claudet et al., 2008). Botsford et al. also suggested protecting between one-third and one-half of a given region, demonstrating that reserves are only effective if they exceed the dispersal range of larvae (Botsford et al., 2001). On land, recent estimates propose that at least 44% of the Earth’s surface should be protected to safeguard biodiversity (Allan et al., 2022), while other calls suggest thresholds as high as 75% (Baillie & Zhang 2018).

### Maintaining food webs: large reserves rather than many small ones

Regarding reserve structure, our study suggests that large, contiguous reserves are more effective than several small, dispersed areas for maintaining biodiversity. At the landscape scale, this configuration consistently promotes both overall diversity and trophic verticality. Reserve aggregation enhances all measures of diversity within protected areas, whereas reserve dispersion primarily benefits local diversity in exploited zones. An ongoing debate in terms of management is whether the disposition of protected area matters (SLOSS debate) or whether the dominant constraint is the total habitat regardless of its configuration (Habitat Amount Hypothesis, Fahrig 2013). Interestingly, our results somewhat reconcile the two views. When considering local properties (eg, alpha diversity), our results suggest, in line with the habitat amount hypothesis, that total protected habitat is the dominant force. However, considering other objectives (gamma diversity, overall verticality, off reserve naturality), the aggregation of reserves certainly matters (in line with the SLOSS debate). This highlights that the definition of objectives and of the scale at which they are measured plays a key role in such debates.

While our results contrast with some empirical findings (*e*.*g*., Fahrig 2020), part of this contrast is linked to our consideration of complex networks (metawebs), a side that is often left neglected or unmodeled in such debates. Our model incorporates trophic interactions and constrains colonisation-extinction dynamics based on the local composition of patches (see Methods). The persistence of top-level consumers depends on the integrity of the underlying trophic chain, which is more likely to be preserved when high-quality patches are spatially aggregated into a larger, continuous protected zone. These results align with the assumptions underlying the ‘minimal patch size requirement’ hypothesis (Hanski, 1994; Lindenmayer et al., 1999) which posits that species require a minimum amount of habitat to persist (Hanski, 1994), and that aggregation facilitates this condition. Moreover, as empirical evidence favoring several small reserves mainly stems from studies focusing on low trophic level species (see references in Fahrig, 2020), explicitly incorporating trophic structure in models can provide novel insights into optimal reserve design. Additional factors may also support the effectiveness of large reserves over smaller ones, particularly when extinction processes dominate colonisation-extinction dynamics (Fahrig et al., 2022). In our model, patches are not necessarily distant from one another (as the results hold across various levels of reserve coverage), but extinction probabilities in exploited areas are consistently higher than colonisation probabilities (see Figure S6). These assumptions reflect the ecological realities of certain systems (Brennan et al., 2022), though they warrant further testing to confirm the robustness of our findings. Moreover, even though such conditions should theoretically favor the “several small” strategy, we still observe the same pattern under higher dispersal scenarios and despite the fact that extinction risks are intrinsically amplified by the presence of antagonistic species (e.g., predators). Lastly, many empirical studies evaluate the SLOSS debate using species accumulation curves (Quinn and Harrison, 1988), which plot the cumulative number of species detected against the cumulative area protected. If, for a given total species richness, the curve formed by accumulating small to large reserves lies above the curve for the inverse order (e.g., concave vs. convex), then the conclusion is that multiple small reserves are preferable. However, our modeling framework allows direct comparison of distinct spatial configurations, each potentially supporting different levels of biodiversity. This avoids inferring the effect of aggregation from a single species accumulation curve, and instead provides a more comprehensive understanding of how reserve structure influences biodiversity retention.

### Improving protection levels

The advised proportion of 30% of protected area proposed by the Kunming-Montreal Global Biodiversity Framework has been taken as a guideline in various countries. It however says little, if the level of protection is left out. While our results suggest that such a level often has limited effects (see discussion above), such effects vastly depend on the protection level (see figure 4). National (or more local) management can be more precise. For instance, in France, the 30% target has been complemented by a “10% of strong protection” target (presumably meaning that the 20 other percent will be less protected). In the light of our results, such levels are not adequate for ecological network protection. Levels of protection in general are very variable. For instance, an analysis of all European marine reserves revealed that exploitation can be even stronger within reserves compared with adjacent areas (Dureuil et al 2018). Our results are consistent with previous calls advocating for the implementation of more effective protection measures within protected areas (Dureuil et al., 2018; Grorud-Colvert et al., 2021; Watson et al., 2014). They also highlight that a strong protection allows to reconcile multiple objectives both inside and outside reserves. Importantly, increasing the level of protection allows for achieving similar conservation outcomes with a reduced total protected area. It also allows higher naturality outside reserves, presumably supporting ecosystem services. Such spillover effects are for instance well studied in the context of marine reserves feeding adjacent exploited areas (Cabral et al 2020, Sala et al 2021). These findings suggest that while setting aside extensive areas may be socially and politically challenging (Kovács et al., 2015), enhancing the quality and effectiveness of protection may represent a pragmatic conservation and conciliation strategy (Dureuil et al., 2018).

### Maintaining diversity in protected and exploited areas

Our study suggests that exploited areas benefit from the presence of protected zones, and that, unlike biodiversity conservation within reserves, the maintenance of naturality outside reserves is enhanced when reserves are spatially dispersed. These findings align with growing calls to develop networks of protected areas that reconcile biodiversity conservation with food production objectives (Gaines et al., 2010). For instance, strategically protecting 28% of the ocean surface could lead to an increase of 6.2 million metric tons in fisheries catch (Sala et al., 2021), largely due to stronger spillover effects from protected to exploited zones (Vandeperre et al., 2011). In terrestrial systems, pest control by natural enemies is more effective in patchy landscapes (see review by Bianchi et al., 2006), in part because increased connectivity enhances spillover processes (Brudvig et al., 2009). Conversely, and in line with our previous results, biodiversity in exploited zones is more negatively affected under land-sharing strategies than under land-sparing ones (Phalan et al., 2011).

### Limitations

Research on the spatial determinants of biodiversity persistence has primarily focused on metapopulations (Hanski and Ovaskainen, 2000; Ovaskainen, 2002) or metacommunities (Mouquet and Loreau, 2003). Network-based approaches have mostly been limited to trophic chains (e.g., Holt, 2002) but demonstrate that their lengths increase with ecosystem size (Holt, 2002), which is consistent with our findings. Our study contributes to this body of literature by exploring these dynamics through a stylized but integrative framework. First, we do not account for competitive interactions (except apparent competition), although dispersal can strongly influence biodiversity in competitive communities (Loreau and Mouquet, 1999), and spatial segregation of reserves might help preserve weak competitors by keeping them away from dominant species (Fahrig et al., 2022). Second, we assumed homogeneous baseline habitat quality across all patches, uniform protection levels in all reserves, and consistent exploitation intensity in non-protected areas. However, habitat heterogeneity is a key factor in shaping biodiversity in spatial food webs (Ryser et al., 2021), empirical studies show that layered protection levels can enhance biodiversity (Loiseau et al., 2021) and variation in exploitation intensity likely plays a critical role in the effectiveness of dispersal. Finally, species introductions or local extinctions can trigger significant bottom-up or top-down cascades at the food web level (Baum and Worm, 2009; Estes and Palmisano, 1974). The strength of these effects may depend on interaction intensity within the network (Barbier and Loreau, 2019), alter effective dispersal rates (Fronhofer et al., 2018), and deserve further exploration to assess their implications for spatial reserve design.

### Openings

Our study advocates for the implementation of large, contiguous reserves to preserve biodiversity and the structural integrity of trophic networks. Although we explored a range of dispersal distances, we assumed a uniform dispersal capacity across all species. In reality, larger species tend to disperse further (Jenkins et al., 2007), which may help explain discrepancies with other studies showing that increased dispersal can reduce local diversity by synchronizing population dynamics (Blasius et al., 1999; Gouhier et al., 2010). Moreover, reserve establishment can select against dispersal at the population level over time (Poethke et al., 2011), but may also buffer species from the evolutionary consequences of exploitation (Miethe et al., 2010). These dynamics could be further explored using our same approach inspired by the *trophic theory of island biogeography* (Gravel et al., 2011) and extended to incorporate evolutionary trade-offs between colonisation and competition capacities (e.g., Tilman, 1994).

Incorporating trophic structure into conservation planning is now essential (Dansereau et al., 2025). Locally tailoring reserve configurations to ecological context, biodiversity hotspots, and food web structure should go hand in hand with efforts to reduce anthropogenic pressure within exploited areas.

## Supplementary materials

**Figure S1:**
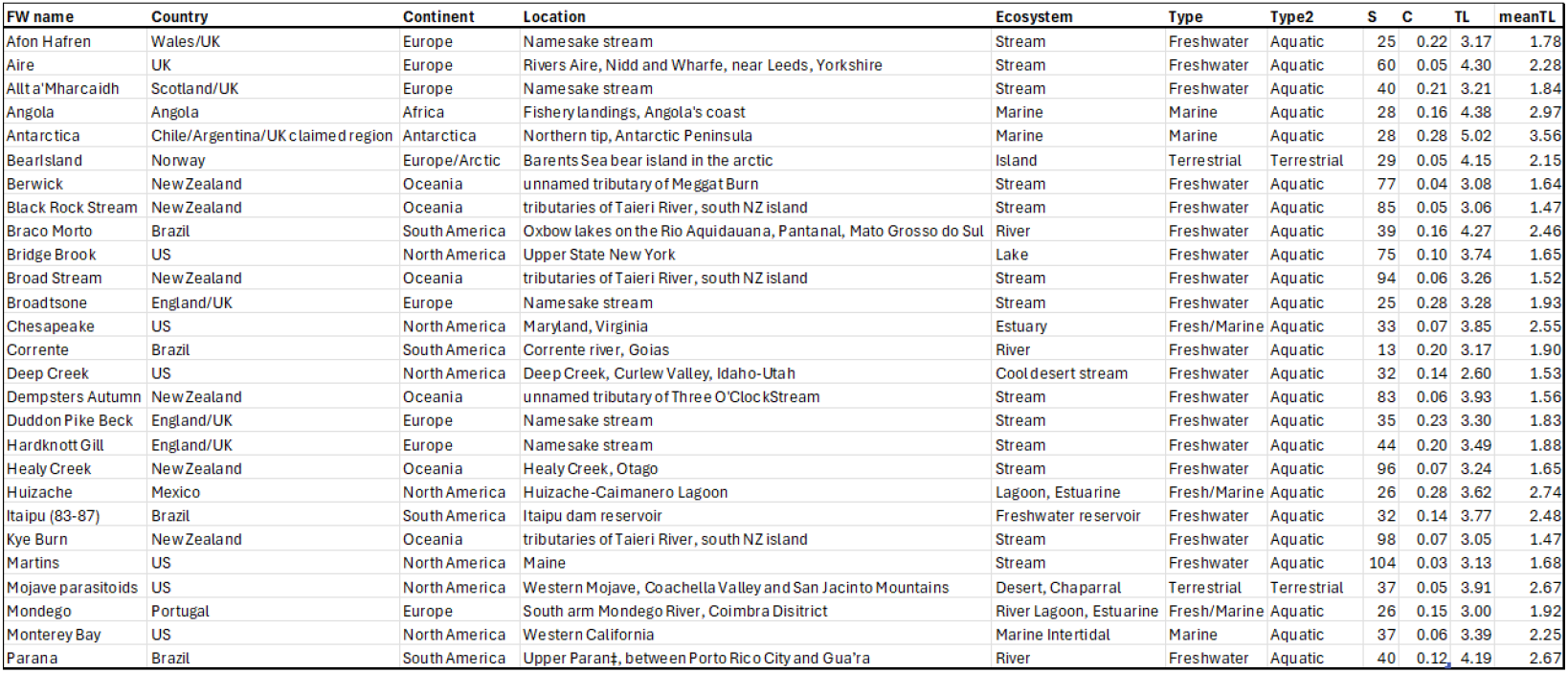
Description of the different networks used in the study.

**Figure S2:**
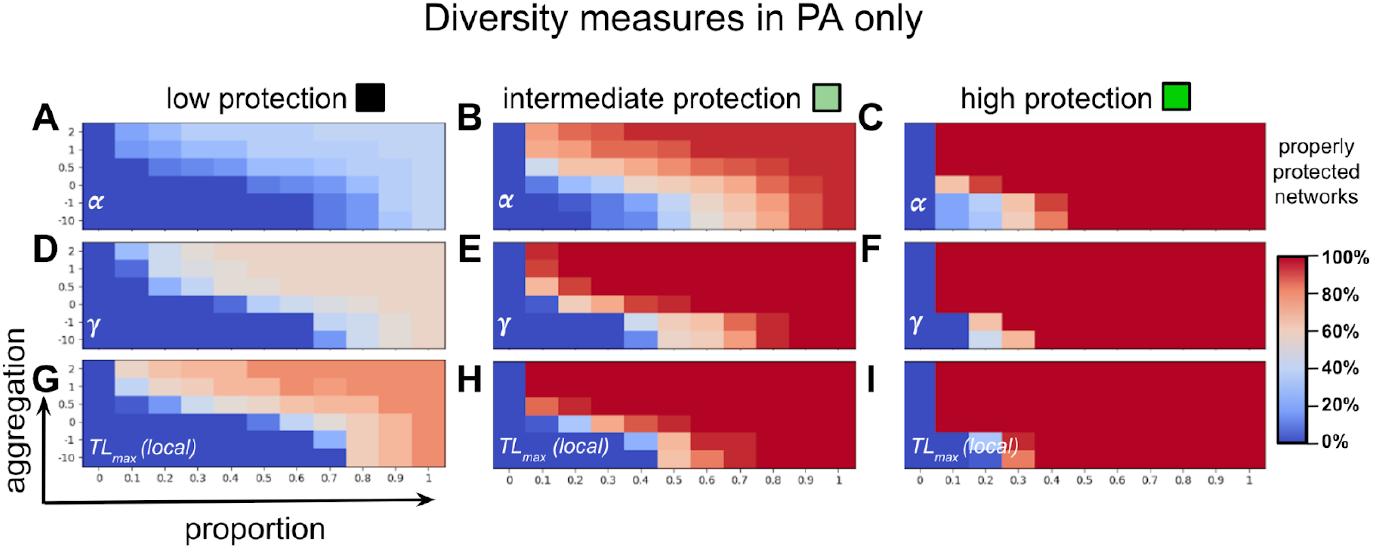
Effectiveness of reserve architecture (aggregation, proportion, and protection level) on achieving conservation targets in protected areas only. A spatial configuration is considered effective for a conservation objective when it maintains the network state above the corresponding threshold. When the network maintains at least 35% of its local diversity (α, A, B, C), it is considered adequately protected. This threshold equals 80% for regional diversity (γ, D, E, F), 50% for the maximum local trophic level (G, H, I). The three columns differ by protection level, which are, from left to right, *p* = 0. 2, *p* = 0. 3 and *p* = 0. 5.

**Figure S3:**
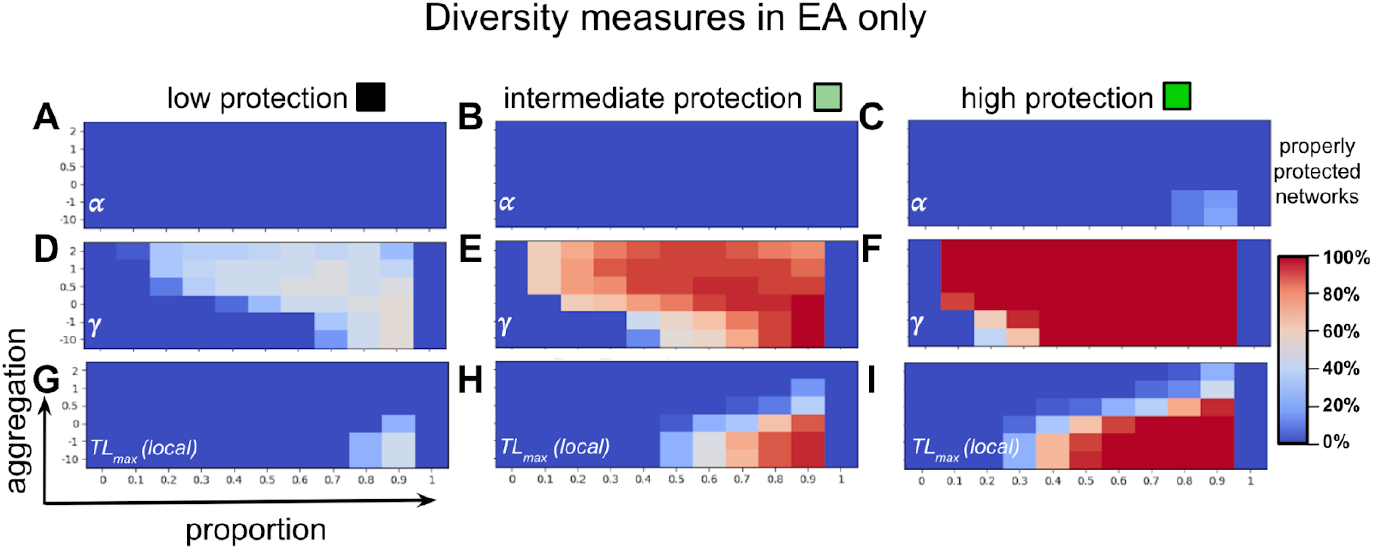
Effectiveness of reserve architecture (aggregation, proportion, and protection level) on achieving conservation targets in exploited areas only. A spatial configuration is considered effective for a conservation objective when it maintains the network state above the corresponding threshold. When the network maintains at least 35% of its local diversity (α, A, B, C), it is considered adequately protected. This threshold equals 80% for regional diversity (γ, D, E, F), 50% for the maximum local trophic level (G, H, I). The three columns differ by protection level, which are, from left to right, *p* = 0. 2, *p* = 0. 3 and *p* = 0. 5.

**Figure S4:**
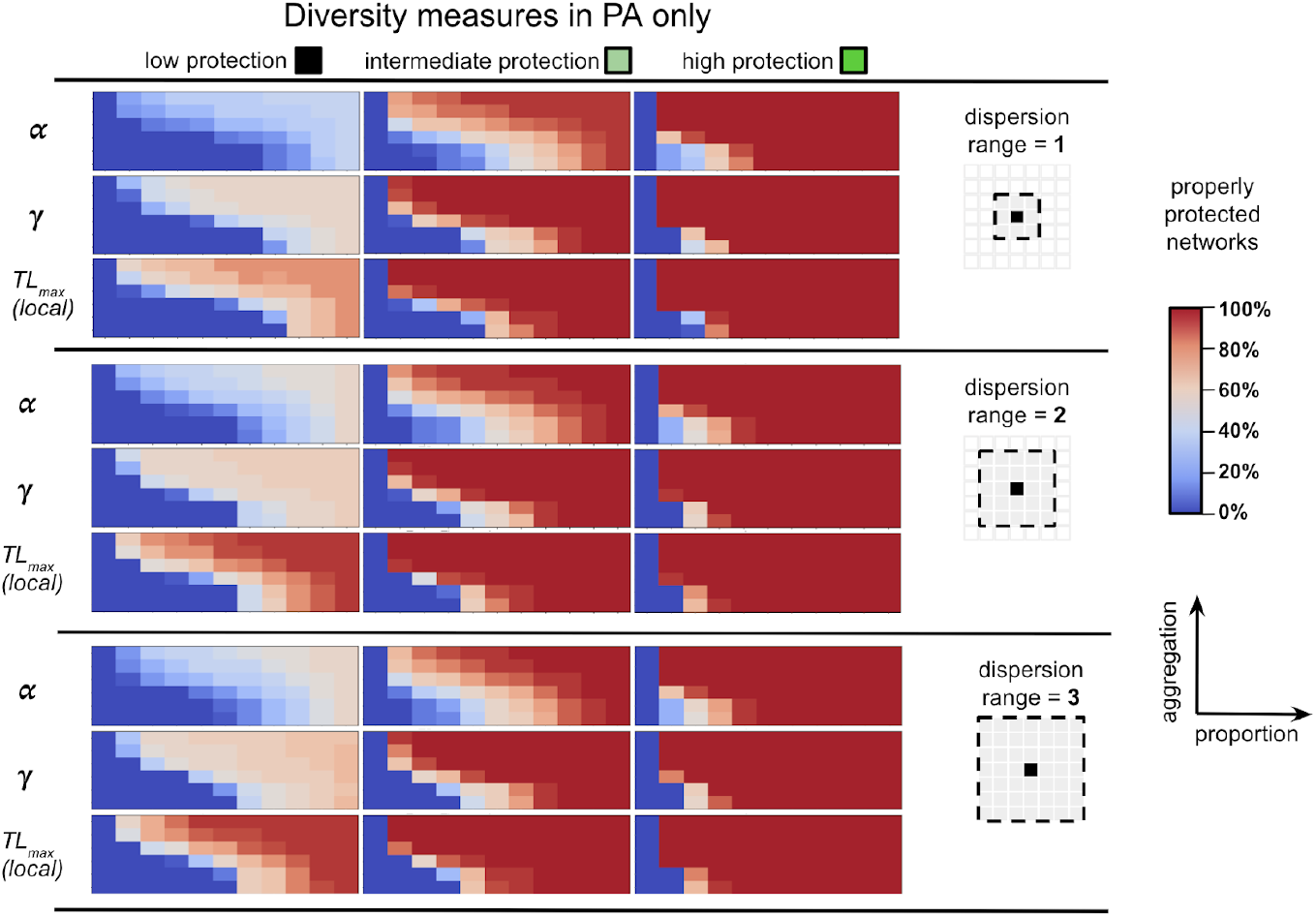
Impact of dispersal range on reserve architecture (aggregation, proportion, and protection level) effectiveness regarding conservation targets in protected areas only. A spatial configuration is considered effective for a conservation objective when it maintains the network state above the corresponding threshold. When the network maintains at least 35% of its local α diversity, it is considered adequately protected. This threshold equals 80% for regional γ diversity and 50% for the maximum local trophic level. The three columns differ by protection level, which are, from left to right, *p* = 0. 2, *p* = 0. 3 and *p* = 0. 5.

**Figure S5:**
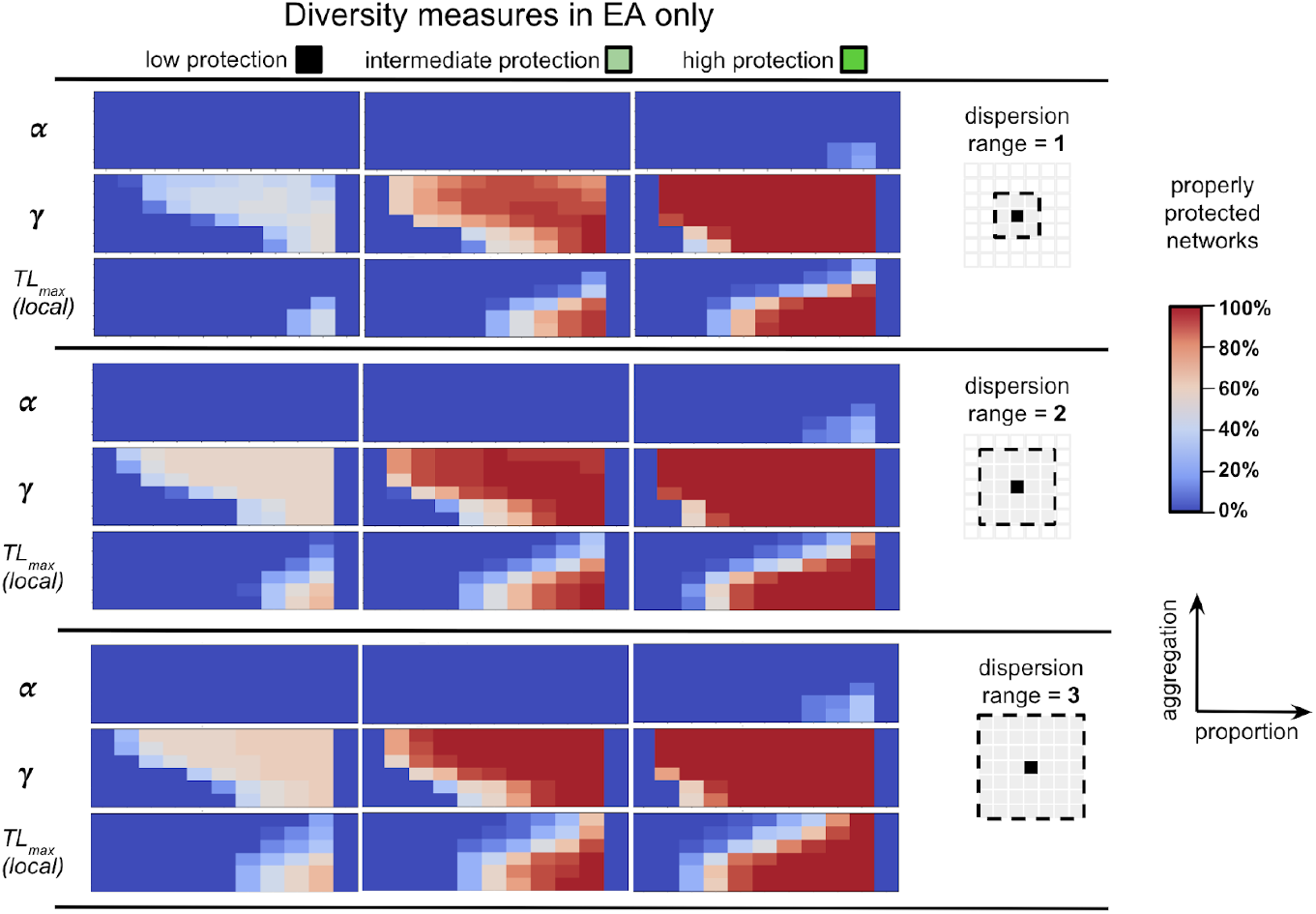
Impact of dispersal range on reserve architecture (aggregation, proportion, and protection level) effectiveness regarding conservation targets in exploited areas only. A spatial configuration is considered effective for a conservation objective when it maintains the network state above the corresponding threshold. When the network maintains at least 35% of its local α diversity, it is considered adequately protected. This threshold equals 80% for regional γ diversity and 50% for the maximum local trophic level. The three columns differ by protection level, which are, from left to right, *p* = 0. 2, *p* = 0. 3 and *p* = 0. 5.

**Figure S6:**
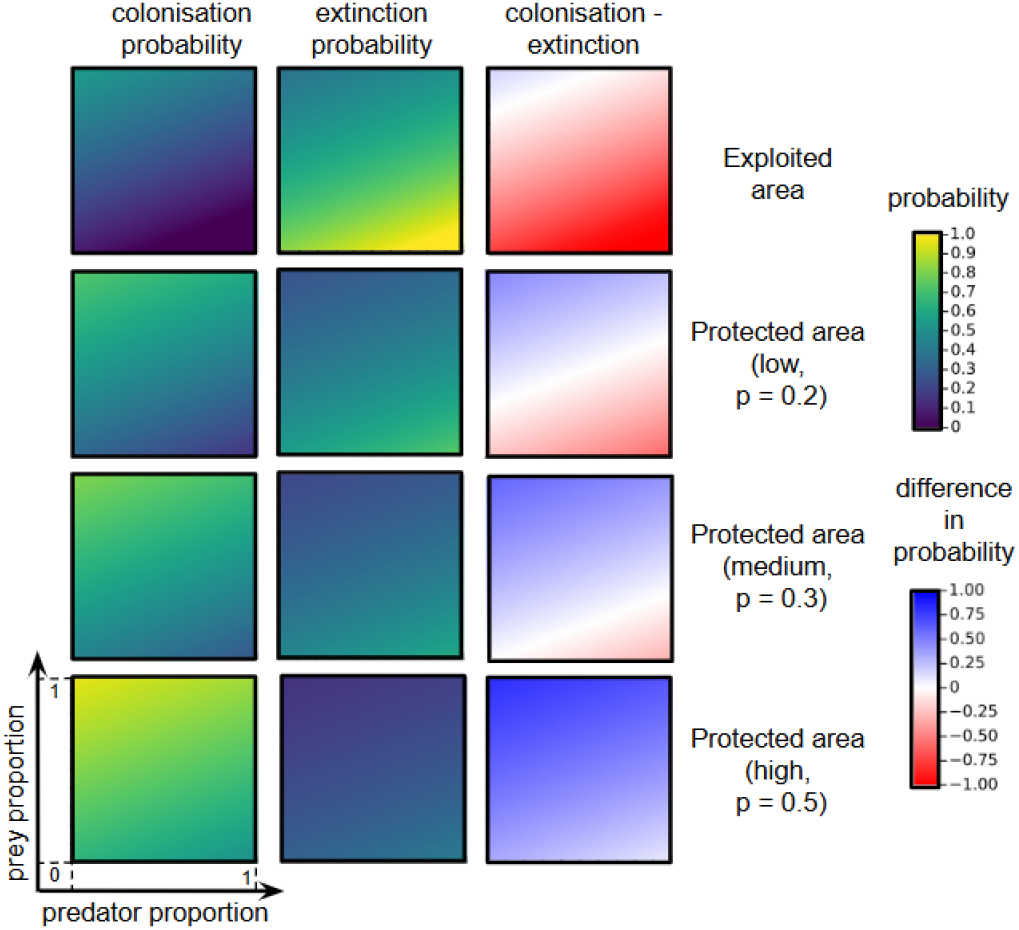
Impact of patch quality on colonisation and extinctions probabilities. Patches with a higher proportion of potential prey species for a given species are more likely to be colonized by that species. Conversely, patches with a higher proportion of potential predators are associated with a higher extinction probability for the focal species. Top panel: exploited zones. Bottom three panels: the different levels of protection within protected areas used in the study. For colonisation probabilities, the focal species is present in all surrounding patches (*n*_*i,j*_ = 1, see eq. 2).

## Notes

### Competing Interest Statement

The authors have declared no competing interest.

### Funding

- Sorbonne Université

